# Robot-Assisted SpiderMass for *in vivo* Real-Time Topography Mass Spectrometry Imaging

**DOI:** 10.1101/2020.12.15.422889

**Authors:** Nina Ogrinc, Alexandre Kruszewski, Paul Chaillou, Philippe Saudemont, Chann Lagadec, Michel Salzet, Christian Duriez, Isabelle Fournier

## Abstract

Mass Spectrometry Imaging (MSI) has shown to bring invaluable information for biological and clinical applications. However, conventional MSI is generally performed *ex vivo* from tissue sections. Here, we develop a novel MS-based method for *in vivo* mass spectrometry imaging. By coupling the SpiderMass technology - that provides *in vivo* minimally invasive analysis – to a robotic arm of high accuracy, we demonstrate that images can be acquired from any surface by moving the laser probe above the surface. By equipping the robotic arm with a sensor, we are also able to both get the topography image of the sample surface and the molecular distribution, and then and plot back the molecular data, directly to the 3D topographical image without the need for image fusion. This is shown for the first time with the 3D topographic MS-Based whole-body imaging of a mouse. Enabling fast in vivo MSI bridged to topography pave the way for surgical applications to excision margins.

---

Mass spectrometry (MS) has, over the past two decades, moved closer to the patient’s bedside. Particularly, the development of Mass spectrometry Imaging (MSI), first with Matrix-Assisted Laser Desorption Ionization (MALDI-MSI) and later with Desorption Electrospray Ionization (DESI-MSI), has spurred interest for molecular imaging in oncology. The technology has demonstrated that collected MS molecular profiles are specific to cell phenotypes and provide access to the molecular content in the tumor microenvironment. On the other hand, the introduction of MS directly in the surgical theater represented a true paradigm shift by transforming MS from a lab technology into a clinical tool. Several intraoperative MS techniques, either for *ex vivo* tissue analysis on the bench, or directly targeting *in vivo,* have been developed. With respect to *in vivo* a few different MS techniques based on direct coupling to either surgical ^1,2^, laser ^3–5^ or extraction devices ^6^ techniques have revolutionized the surgico-clinical field. These *in vivo* techniques target the same goal by helping practitioners with decision making during surgery. To date *in vivo* MS has only been demonstrated by manual analysis of defined spots on the tissue, and for classification purposes of molecular profiles. However, to provide the global “picture” to the surgeon for improved decision making, a complete area needs to be imaged by an autonomous system. To achieve *in vivo* MS Imaging several constraints need to be taken into consideration including that i) the sample cannot be moved under the analytical beam and ii) the biological tissue must not be altered in any way. Indeed, conventional MSI is realized from flat samples surfaces with the sample probe moving under the analytical beam. Some instruments such as the TOF-SIMS and a few MALDI systems offer the possibility to electrostatically move the analytical beam at the scale of the field view but only for a very limited area (<0.1 mm^2^). A way to conduct *in vivo* imaging is to develop a system where the analytical beam is moving in a well-controlled fashion in 3 dimensions with enough accuracy above the sample surface to address rugged surfaces.

Here, we introduce topography molecular imaging by robot-assisted MSI as a novel way to achieve *in vivo* imaging. The topography MSI is designed to image raw samples in real-time while providing both the topography and the related molecular information. We developed a robotic arm system coupled to the Water-Assisted Laser Desorption Ionization technology (SpiderMass). SpiderMass is an emerging mini-invasive IR laser-based MS technique designed for *in vivo* real-time molecular analysis^3,4^ and has been showcase at the veterinary surgery room for tissue profiling^7^. The method is based on the resonant excitation of the endogenous water molecules and provides metabo-lipidomic molecular profiles specific to different cell phenotypes very similarly to the others MSI modalities (e.g. MALDI or DESI)^7^. One of the major advantages of the SpiderMass is the ability to manually move the probe and freely scan any surface without constraint or contact to the specimen. However, the system as such does not allow MSI to be performed. The development of a combined system including a robotic arm enables, thus, the automated fine motion with the requested precision of the laser microprobe above the sample surface for image acquisition. We also demonstrate here, that by integrating a sensor to the system which can provide height measurement, a topographic image of the surface can be obtained. With this system, topography MSI can be achieved from biological sample with a spatial resolution of 400-500 μm both for the topographical measurement and the molecular MSI.

## RESULTS

### Topography MSI design and setup

We started with designing and setting up the system with the perspective to realize *in vivo* MSI in humans in the future. The careful design and developments were executed in 3 different modules including i) hardware adjustments, ii) an automatized user interface, and iii) image reconstruction. The hardware is presented in **Fig. 1**. It includes a 6-axis robotic arm with 5 μm repeatability and 100 μm path accuracy, and an adaptor equipped with a LED and a camera cooperating as a distance sensor. The SpiderMass microprobe is mounted on the arm through the adaptor, including the laser fiber and the transfer line connected to the MS instrument (**Fig. 1a, 1b**). The zoom into the adaptor specifies the placement of the camera and the LED. The arrows indicate the possible motions of the motors. The photos of the real system setup are shown in **Fig 1b**. The general workflow of a 3D imaging experiment includes 3 steps: 1/ the topography recording, 2/ collection of the molecular data and 3/ the image reconstruction (**Fig. 1c**). The images are acquired within the first two steps. First, to start the acquisition, the distance sensor is calibrated by measuring the distance between the LED and the center of the observed camera field. Then the investigated specimen is placed under the arm and the (X, Y, Z) coordinates are recorded point by point of the defined imaging area to create the topographical map. To accelerate this step, the resolution of the map can be lowered to reduce the number of collected data points. An interpolation is used to reconstruct the surface at required spatial resolution and stored. Second, the topography is used to correctly position the laser probe. Several shots are fired in a burst at each data point while the molecular information is collected by the mass spectrometer. The raw data files are converted into the vendor format as an mzXML file for data extraction and normalization. Finally, the *m/z* values of interest are selected, and the images are reconstructed and displayed.

**Fig. 1.**
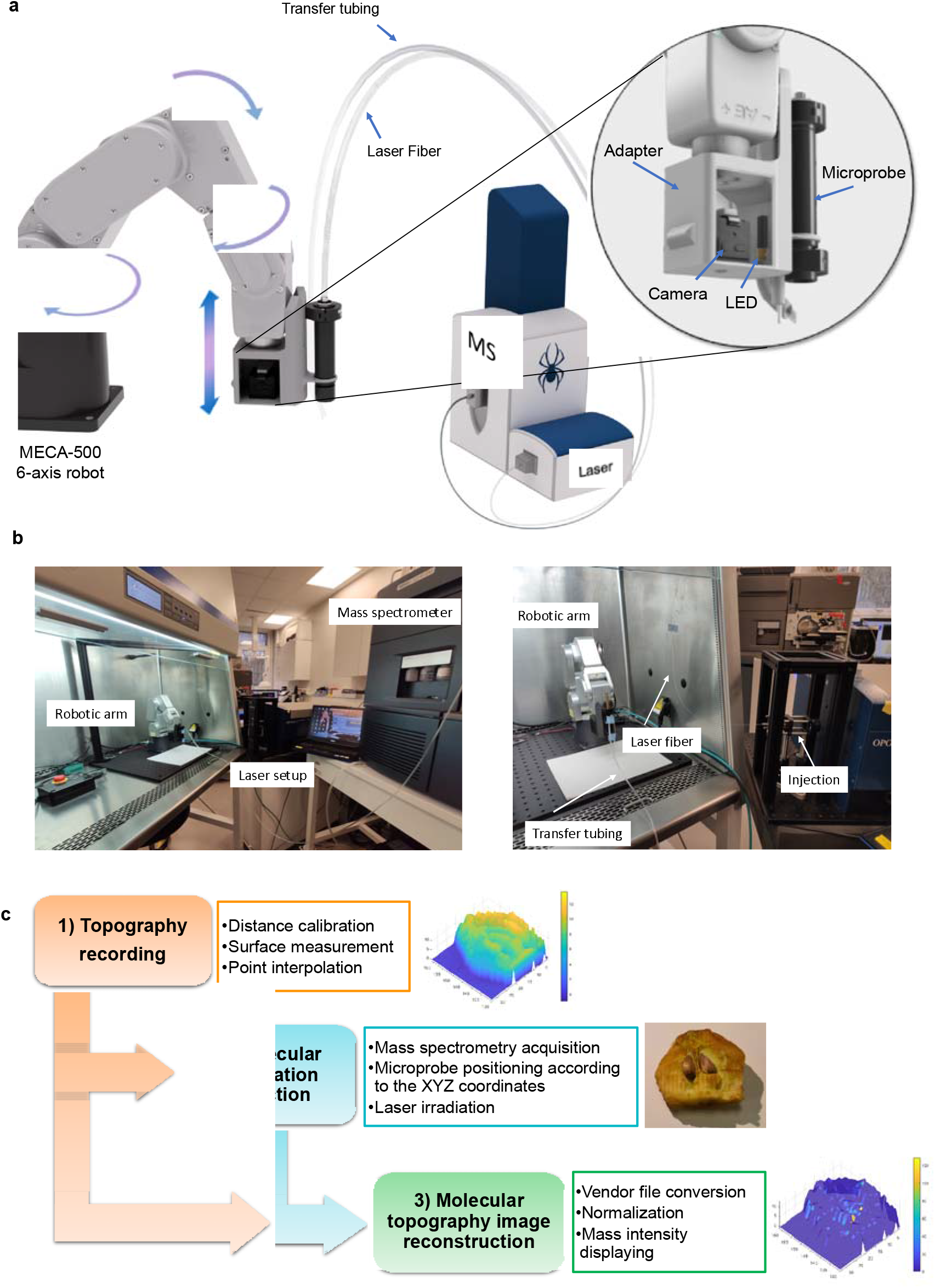
Schematic representation of the Robot-Assisted SpiderMass technology and the general workflow of operation. **a**, The SpiderMass probe is attached with an adaptor to the robotic arm. The zoomed part details the distance sensor system with a separate compartment for the camera and an LED laser pointer. The arrows indicate some degrees of movements of the 6-axis robotic arm. The robotic arm manipulates the laser microprobe i.e. the handpiece placed at the exit of the optical fiber and the transfer tubing where the ablated material is sent by aspiration back to the MS instrument. **b** Photos of the robot-assisted SpiderMass MSI imaging setup. The layout of the whole system including the robotic arm, laser system and the mass spectrometer (left) and the focus on the robotic arm with the laser fiber and transfer tubing positioned in the fumehood and the laser injection (right). **c** Three-step general workflow of a 3D imaging experiment. First the x, y, z topography is recorded and stored for the correct laser positioning, then the molecular information is collected by a mass spectrometric acquisition, and finally the image is reconstructed and displayed.

### Topographical MSI of model samples

We tested the automated robotic arm on several model objects and biological samples starting with the topography imaging step. Examples of topographical maps indicating the objects native shapes and forms are shown in **Fig. 2**. To determine the accuracy of the z-height profiles we created topographical maps of a USB key (**Fig. 2a**), an apple core with seeds (**Fig. 2b**) and the surface of a small piece of sponge (**Fig. 2c**). Each map includes a color bar indicating the height differences (yellow= high /blue =low). The USB key and apple core piece show the total difference in height to be between 0-12 mm, while the z height of the sponge surface varies between 14 and 18.3 mm which are perfectly in line with the topography of these samples. In a second step the molecular MS-based imaging is realized. The SpiderMass laser microprobe is re-scanning the specimen, following the same starting position and topography, including z-height to retain the microprobe in the correct focal plane.

**Fig. 2.**
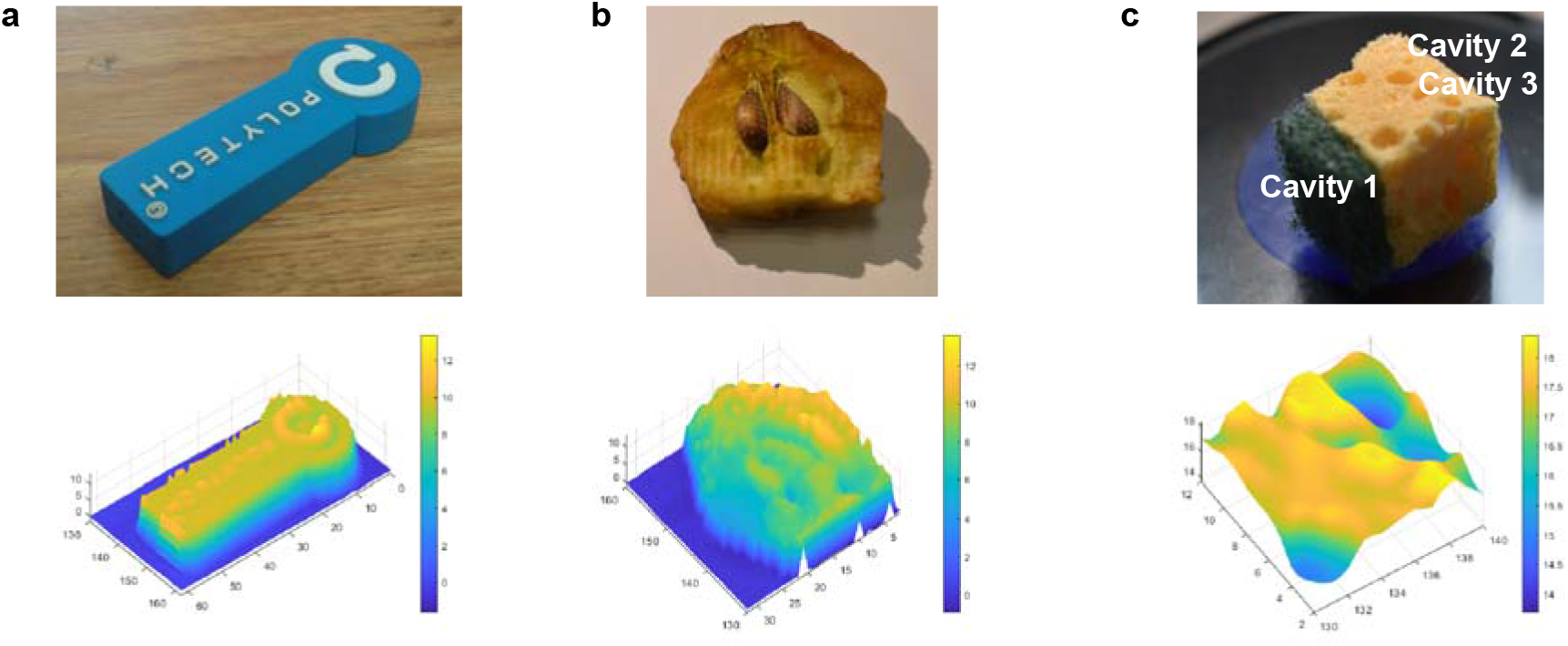
Topographical images created using the topography MSI. Optical and topography maps of **a** USB key, **b** apple core with seeds and **c** 1 cm^2^ surface of a sponge piece. The color bars indicate the height differences in mm.

Following the 2-step scanning, the topographical and molecular images were coregistered for the small piece of sponge (**Fig. 3**, upper panel). In this example, an imaging area of 1cm^2^ was selected on the surface of the sponge and scanned with a 500 μm step size both for the topography (**Fig. 3a**) and the MS Imaging (laser beam diameter at focal point 400 μm). The topographical image (**Fig. 3b**) shows the z-height difference on the imaged surface to be 4 mm with the deepest z at the cavities. The 3 most pronounced cavities in the topographical map are indicated in the optical image (**Fig. 2c**). The first cavity is 2mm in depth as indicated by the topographical image and confirmed by measuring with a ruler **(Supplementary Fig. 1)**. The total acquisition time of the molecular imaging by SpiderMass was 24 min. The spectra were recorded in positive ion mode and showed good signal intensity with the most predominant peaks from the sponge detected at *m/z* 372.3, 575.3 and 675.6 (**Supplementary Fig.2**). Because the composition of the sponge is unknown, a second acquisition was performed the of the same area with the addition of a 1mL of a lipid standard mix solution containing [PC(34:1)+H]^+^ and [PC(32:1)+H]^+^ inside of the cavities. The Total Ion Current (TIC) chromatogram and first indicative pixels are shown in **Supplementary Fig. 3**. Each chromatographic peak corresponds (red bar) to one pixel on the topographical map. A group of chromatographic peaks (blue bar) represent on line in the topographical image (**Supplementary Fig 3**). The TIC absolute intensities of the spectra were increased to 1e^7^ after spotting the standards and new signals are observed in the *m/z* [700-900] range (**Fig. 3c**) including *m/z* 732.6 ± 0.1 and *m/z* 760.6 ± 0.1 corresponding to the two lipid standards. These signals were used for the MSI reconstruction and plotted to the topographical image (**Fig 3d** and **e**), respectively. As indicated in the images, the highest normalized intensities of the two lipid peaks are originating from within the cavities where the lipid standards were deposited.

**Fig. 3.**
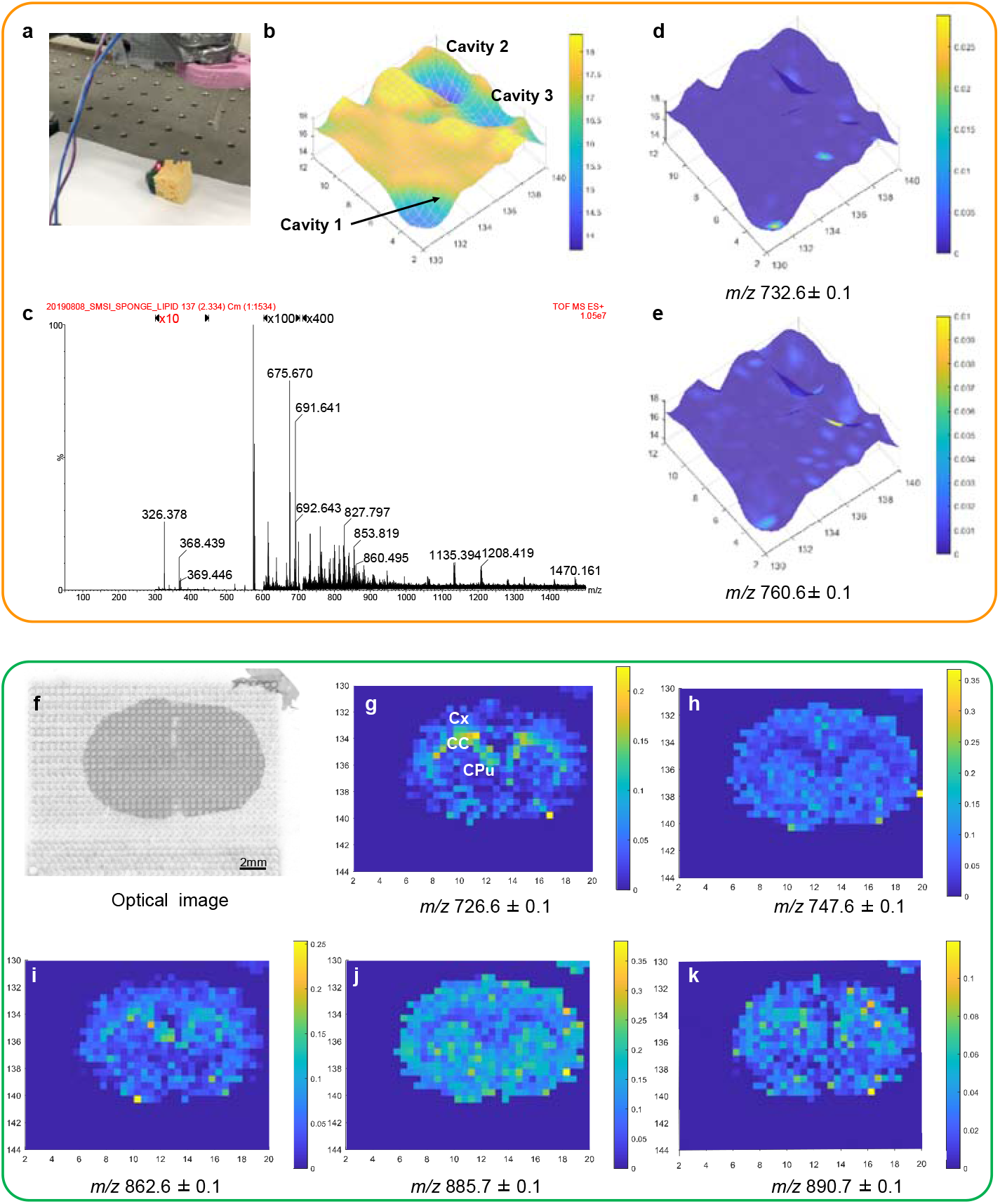
Robot-Assisted SpiderMass MSI on model samples. Images of the sponge with cavities and rat brain tissue section. **a**Photo of the topographical scanning with the distance sensor to create the topographical image. **b** Topographical 1cm^2^ mesh of the sponge with indicated cavities. The color bar indicated the topographical changes in the z dimension. **c** Total Ion count mass spectrum obtained from the sponge surface following the topographical mesh**. d-e** The plotted m/z values in positive ion mode on the topographical image with normalized intensities (732.6±0.1 and 760.6±0.1). Color bars indicate higher intensity in yellow and lower intensities in blue. **f** The optical image of the rat brain tissue following the Robot-Assisted acquisition. **g-k** Plotted m/z values in negative ion mode on a 1.8 cm × 1.4 cm 2D map of rat brain clearly indicating the tissue outline with some distinctive regions. The ion images correspond to *m/z* 726.6 ±0.1 PE (18:1/18:1)-H]^-^, 747.6 ± 0.1 [PA (22:6/18:0)-H]. *m/z* 862.6 ± 0.1 SHexCer 18:1;2_22:0/ [PC(P-18:0_22:1))+Cl]. *m/z* 885.7± 0.1 [PI(18:0/20:4)-H]^-^, and *m/z* 890.7 ± 0.1 SHexCer d18:1/24:0 / [PC(O-42:2))+Cl]^-^ (b-f) with TIC normalized intensities. The color bars indicate higher intensity in yellow and lower intensities in blue. Abbreviations: Cx - cerebral cortex; CC - corpus callosum; CPu - caudate putamen.

The user interface was developed to also enable the acquisition and visualization of 2D images from tissue sections (**Figure 3**, bottom panel). The applicability of the 2Dimaging mode was tested on a rat brain histological tissue section. First, a virtual topographical image was created with requested dimensions and then the MSI acquisition was conducted following the same pattern in the negative ion mode (see **Figure 3f**, optical image). The images were co-registered in the same manner as previously described for the topographical molecular images. The acquired TIC mass spectrum in the negative ion mode is shown in **Supplementary Fig. S4**. Some of the most prominent peaks in the lipid *m/z* [700-1000] range were further selected for coregistration of the molecular and topography images.; *m/z* 726.6 ± 0.1 [PE (18:1_18:1)-H]^-^, 747.6 ± 0.1 [PA (22:6_18:0)-H]^-^, *m/z* 862.6± 0.1 SHexCer 18:1;2/22:0/ [PC(P-18:0_22:1))+Cl]^-^, *m/z* 885.6±0.1 [PI(18:0_20:4)-H]^-^ and *m/z* 890.7 ± 0.1 SHexCer d18:1/24:0 / [PC(O-42:2))+Cl]^-^, respectively (**Fig 3 f-k**). The images reveal a clear depiction of the white and gray matter in the rat brain tissue. Particularly, ions at *m/z* 726.6, *m/z* 862.6 and *m/z* 809.7 are present in the Corpus Callosum (CC) while the ions at *m/z* 747.6 and *m/z* 885.7 are present across the whole tissue section and depleted in the CC. The MS/MS identification of the lipid species can be found in **Supplementary Table S1**.

### 3D topographical molecular imaging of biological specimens

We further investigated 3D topographical molecular imaging from analyzing several biological specimens namely an apple core with seeds, a skin biopsy disk and within the whole body of a mouse model. In the first example, topographical and molecular images of the apple core with seeds were recorded (**Fig. 4**, upper panel). The apple cross-section consists of the mesocarp, endocarp, seeds and loculus region (**Fig. 4a**). First, the 3D map was created with 3D and 2D view (Fig. 4b and 4c, respectively) revealing different apple core regions. The mesocarp, endocarp and seeds show a higher topography than the loculus region. Several ions were reconstructed on the 3D map (**Fig. 4 d-f**). Specific ions such as *m/z* 664.6 ± 0.1 and *m/z* 700.6 ± 0.1 were more abundant in the seeds and endocarp, while *m/z* 737.6 ± 0.1 was more abundant in the mesocarp. The putative IDs can be found in **Supplementary Table S2**.

**Fig. 4.**
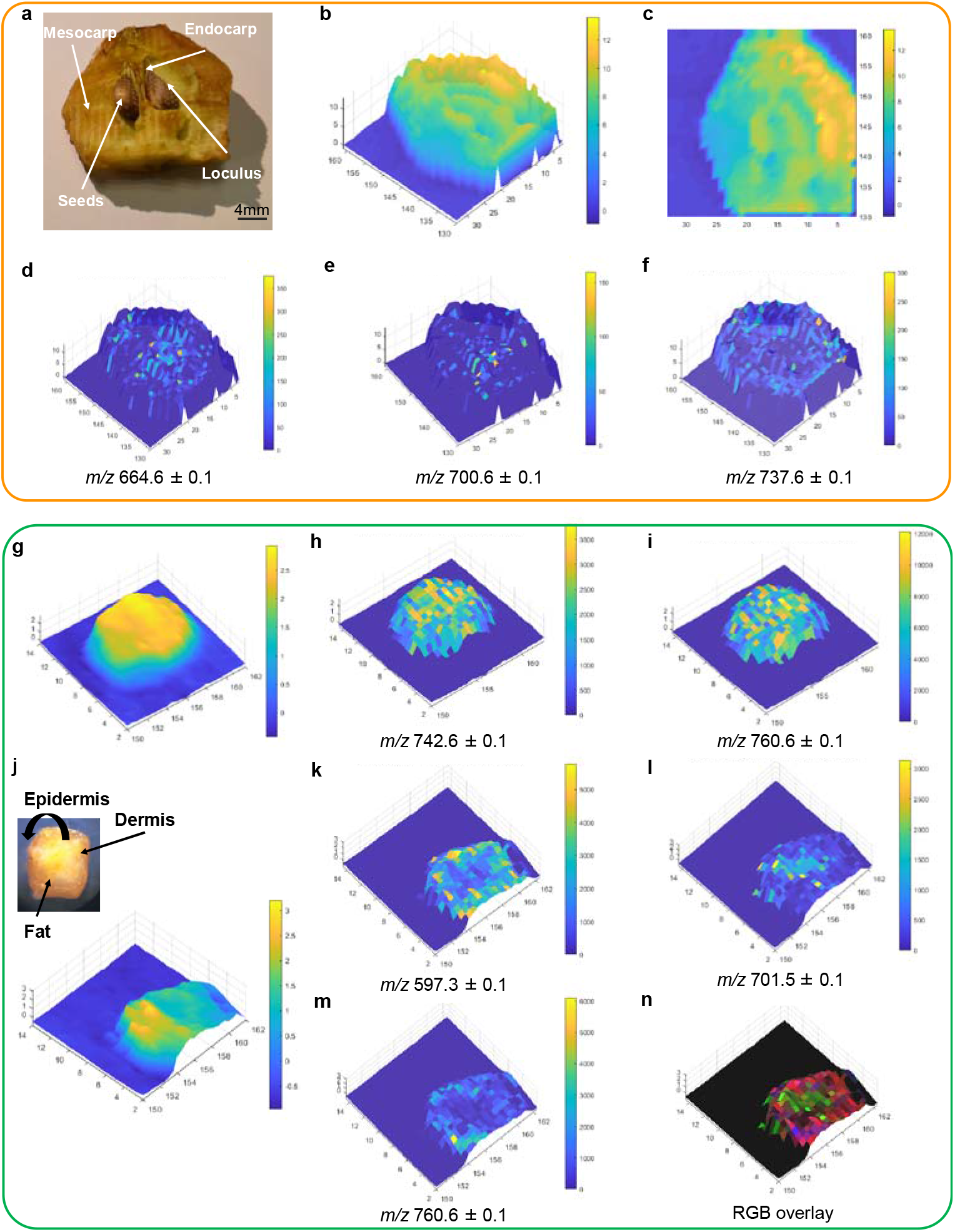
Robot-Assisted SpiderMass mass spectrometry images of biological samples. Images of the apple core with seeds **a-f** and fresh skin disc biopsy **g-n. a** The optical image of the apple crosssection with different regions. The 3D topographical map of the apple core in 3D **b** and 2D mode **c** with color bars expressing different heights. The plotted m/z values clearly showing distinctive regions and correspond to **d** m/z 664.6 ± 0.1, **e** 700.6 ± 0.1, and **f** 737.6± 0.1. Topographical images and an optical image of the fresh skin biopsy acquired from the **g** top and **j** bottom with annotated regions. **h-i** Selected ion images from the skin biopsy acquired from the top at *m/z* 742.6 ± 0.1 and *m/z* 760.6 ± 0.1. **k-n** selected ion images and RGB overlay of the fresh skin biopsy acquired from the bottom at *m/z* 597.3± 0.1 red, *m/z* 701.6 ± 0.1 green and *m/z* 760.6 ± 0.1 blue. All the selected ion images were normalized to the TIC.

We next imaged a fresh abdominal skin biopsy (**Figure 4**, Bottom panel). The skin biopsy is composed of the epidermis, dermis and fatty tissue. Since all different layers cannot be imaged in a single experiment, we conducted experiments from the top (**Fig. 4g**) and bottom (**Fig. 4j**) of the biopsy. The epidermis layer is relatively homogeneous (**Fig. 4g**); hence no spatial molecular differences were observed as corroborated by the distribution of two ions at *m/z* 742.6 ± 0.1 (**Fig. 4h**) and 760.6 ± 0.1 (**Fig. 4i**). In contrast, clear molecular differences were observed between the dermis and the fatty regions acquired from the bottom part of the biopsy (**Fig. 4k-n**). The distribution of the ions at *m/z* 597.3 ± 0.1 and *m/z* 760.6 ± 0.1 is mainly found in the dermis region (**Fig. 4e**), while the *m/z* 701.6 ± 0.1 is mainly distributed in the fat (**Fig. 4f**). The RGB overlay of the selected ions is displayed in **Fig. 4n** (red, *m/z* 597.3 ± 0.1; green, *m/z* 701.6 ± 0.1, and blue, *m/z* 760.6 ± 0.1) and shows the different distribution on the ions in the bottom part of the biopsy. The MS/MS identification of the lipid species can be found in **Supplementary Table S3**.

To explore the outmost potential of the technique we conducted experiments inside of the whole body of a freshly sacrificed mouse. The whole-body mouse optical and topographical images are shown in **Fig. 5a-c**. The images clearly depict different organs such as the intestine, stomach, liver, lungs, and heart. For molecular corelation several smaller areas were imaged including the mammary gland, the heart and the lungs, and brain (**Fig. 5d,g,j**). The collection of topographical images and molecular data are shown in **Video 1.**The first sequence displays the robotic arm setup and the acquisition of the topographical image of the mammary gland. Once the topographical image was collected, the SpiderMass probe followed the topographical pattern in sequence two. All of the selected areas in **Fig 5** were imaged in the same manner. The third sequence depicts the collection of the molecular data from the brain. The video clearly shows the SpiderMass laser spot size (500 μm) and the path of the microprobe following the topographical image. **Fig. 5 e-f** depicts the topographical and molecular images of the mammary gland. While the topographical image reveals only slight differences (2.5 mm, **Fig. 5e**) between the mammary gland and the subcutaneous tissue we can clearly distinguish them molecularly **Fig. 5f**. Several ions, such as *m/z* 630.6 ± 0.1 and *m/z* 734.6 ± 0.1 [PC (32:0) + H]^+^ show higher abundance in subcutaneous tissue, while the ion *m/z* 876.8 ± 0.1 is present only in the mammary gland. This is even further exemplified by the RGB image of the selected species (red, *m/z* 876.8± 0.1; green, *m/z* 734.6± 0.1 and blue, *m/z* 630.6 ± 0.1). These examples demonstrate that molecular data can be plotted back onto topological surfaces of organs bringing molecular knowledge *in vivo* that could be used for further learning or molecular identification. In addition, the mean positive ion mode spectra of the mouse imaging data sets (**Fig. 6a** mammary gland, **Fig. 6b** heart and lungs and **Fig. 6c** brain) show good sensitivity (in the range 1.0e^6^ at the 600-1500 mass range) with hundreds of different lipid species observed. In the more complex region, such as the heart, we notice greater topographical differences **Fig. 5h**. There is a clear variation in height between the heart and lung and the rest of the imaged area. Some selected ions were plotted back to the topographical image in **Fig. 5i**. For example, ion *m/z* 616.2 ± 0.1 corresponding to heme and most probably associated to hemoglobin is distributed mainly in the heart and lungs, while *m/z* 734.6 ± 0.1 [PC (32:0) + H]^+^ is mainly distributed in the lung and surrounding area of the heart. Several ions in the higher mass range such as *m/z* 844.6 ± 0.1 depict the fatty connective tissue surrounding the heart. These tissue specific differences are even more pronounced in the RGB overlay image (red, *m/z* 616.2 ± 0.1; green, *m/z* 734.6 ± 0.1 and blue *m/z* 844.6 ± 0.1). In the last example, the brain images (**Fig. 5k,l**) reveal the expected structure separating the right and the left hemisphere. Some detailed topography of the scalp, midbrain, cerebellum is also found in **Fig. 5k**. Some selected ions are plotted to the topographical image in **Fig. 5l**. The pronounced heme ion at *m/z* 616.2 ± 0.1 shows the distribution across the whole brain imaged area. This is also consolidated by the optical image displaying the imaged zone **Fig. 5j**. The ion *m/z* 734.6 ± 0.1 corresponding to [PC (32:0) + H]^+^ is mainly distributed in the left hemisphere and the cerebellum. This is of no surprise since the particular lipid specie has been repeatably found in the gray matter of the brain. Although we would expect the distribution of this ion to be the same in the right hemisphere there appears to be a slight difference. This can also be observed in the optical image (**Fig. 5j**), where a part of the right hemisphere doesn’t appear to be imaged probably because of the dissection when the skull had to opened. A unique distribution is unraveled by the *m/z* 834.6 ± 0.1 [PC (40:6)+H]^+^ of the surrounding scalp and cerebellum. The RGB overlay depicts the exact features of the brain (red, *m/z* 616.2 ± 0.1; green, *m/z* 734.6 ± 0.1 and blue, *m/z* 834.6 ± 0.1) as shown in the optical image. The MS/MS identification of the heme and lipid species can be found in **Supplementary Table S4**.

**Fig. 5.**
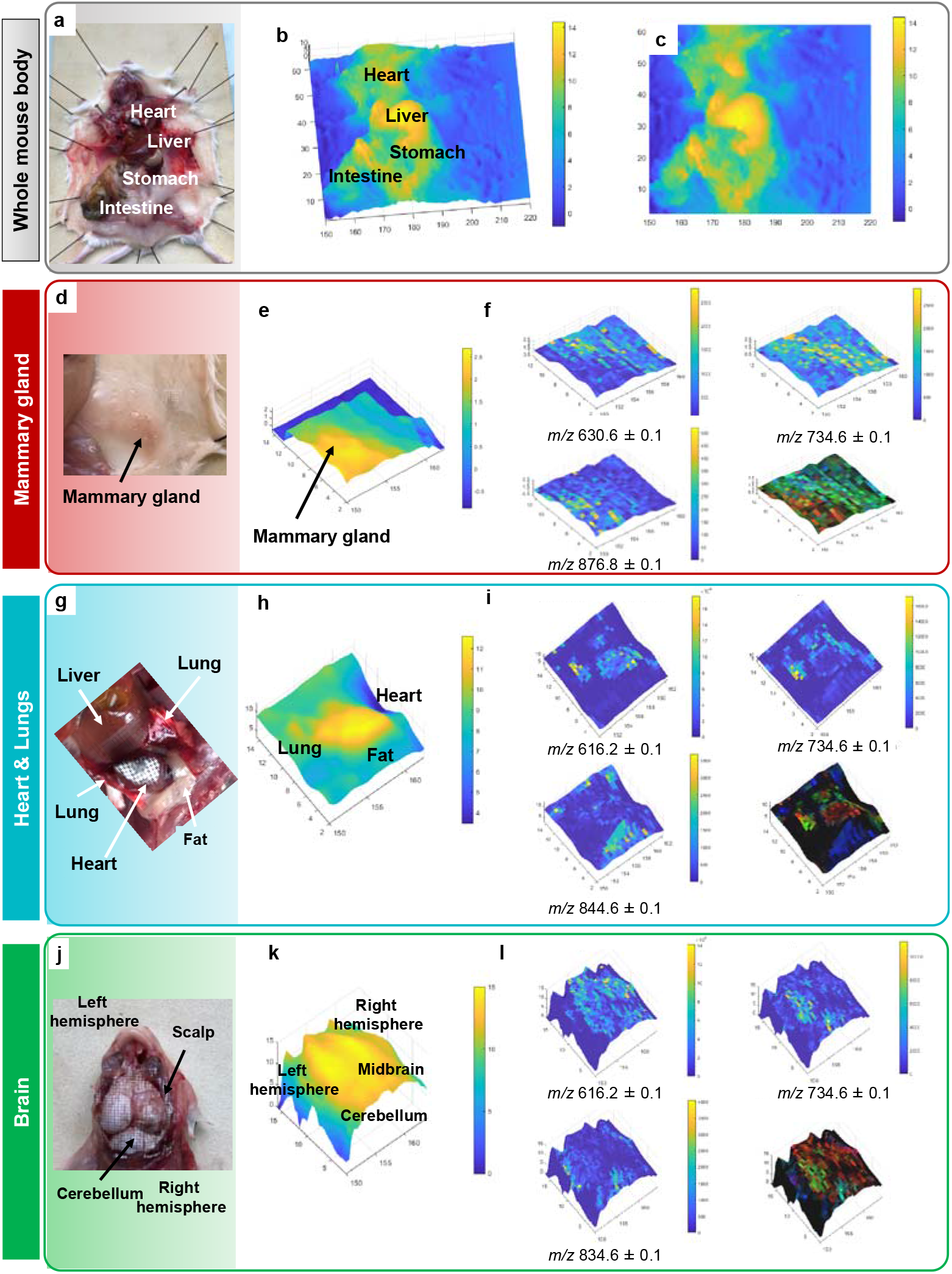
Post-mortem mouse imaging experiments. Optical images of the **a** whole mouse and selected imaging zones; **d** mammary gland, **g** heart and lungs, and **j** brain. Topographical images of the whole mouse in the **b** 3D view, and **c** 2D view. Topographical images of the selected zones depicting different organs and sub-regions; **e** mammary gland **h** heart and lungs, and **k** brain. The color bars indicate the different topographical heights within the imaged area. **f** selected ion images and RGB overlay for reconstruction red *m/z* 876.8; green *m/z* 734.6, blue *m/z* 630.6 of the mammary gland. **i** selected ion images and RGB overlay for reconstruction red *m/z* 616.2; green *m/z* 734.6, blue *m/z* 844.6 of heart and lungs. **l** selected ion images and RGB overlay for reconstruction red *m/z* 616.2; green *m/z* 734.6, blue *m/z* 834.6 of the brain.

**Fig 6.**
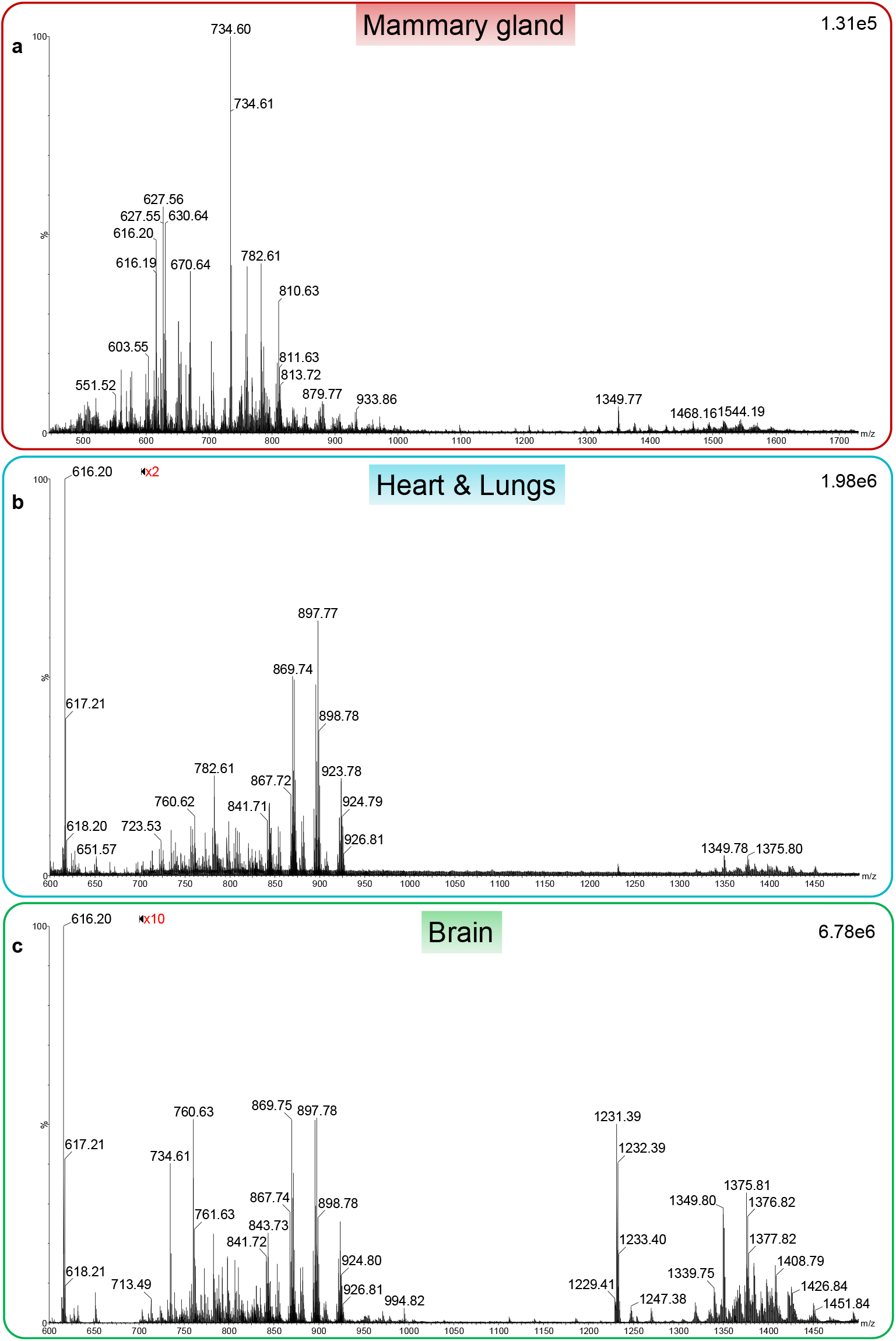
Mass spectra of selected organs from mouse imaging experiments. The representative positive ion mode spectra in the 600-1500 mass range of **a** mammary gland, **b** Heart and lungs, and **c** brain. The color bars indicate the selected ions for molecular image reconstruction and RGB overlays. The absolute intensities for the mean spectra at the selected mass range are indicated in the top right corner and reach over 1e^6^. The spectra are scaled to the highest peak of the selected mass range.

## DISCUSSION

Here we have shown the foundation for the development of *in vivo* real time topographical molecular imaging by the SpiderMass technology. We have successfully coupled the SpiderMass to a stiff 6-axis robotic arm and for the first time demonstrate the possibility to conduct MS Imaging by moving the probe through the assistance of a robotic arm and co-register the molecular information directly on the topographical image matrix pixel by pixel at 400-500 μm resolution without the need for image fusion of the two different modalities. Very few MS devices have been coupled to robotic or automated systems. For example, robotic surface analysis using a needle sampling probe attached to the robotic arm was previously described^8^. Using a 3D laser scanner, Li *et al* created the 3D images of an object surface and analyzed single spots (0.2mm × 1.0mm × 0.6 mm) on the surface by MS using the robotic controlled injection in an open port sampling interface. The molecular information of the analyzed spots was than plotted back to the 3D image but no pixel by pixel co-registration was obtained during the experiments. Collection and data registration are the most crucial parts of imaging experiments. Image fusion is the most common form of co-registration of MSI images in a multimodal approach^9–12^. Here, we plot the distribution of intensities at selected m/z directly on the topographical images via user defined software. Through interpolation, the real time topographical scanning can be achieved within minutes at the desired spatial resolution. The current images were acquired at 400-500 μm resolution to demonstrate the potential of the technique. In the future, we plan to adapt the system to create higher spatial resolution images (~100 μm) by improving the focusing device of the SpiderMass microprobe. However, even at the yet used resolution we were able to topographically and molecularly differentiate certain areas of the brain tissue, apple core and seeds, skin punch biopsy as well as areas within the mouse body. The selected ions from the rat brain tissue section have already been previously detected and identified with other mass spectrometry imaging techniques such as MALDI-MSI or LAESI-MSI^13^. More concretely we were able to discriminate specific lipidomic features corresponding to different tissue types of the skin punch biopsy, mammary gland, subcutaneous tissue, heart, lung and brain. Although the heme ion *m/z* 616.2 is very evenly distributed in several organs as expected, several lipid species such as [PC (32:0) + H]^+^ at *m/z* 734.6 were found specific for lung tissue, subcutaneous tissue and brain tissue. The technology, therefore, clearly demonstrates the ability to discriminate tissue specific features through topographical and molecular images and can be further used to discriminate regions such as resection margins or different cancer subtypes which will help during surgical evaluation.

We have shown the ability to create 2D and 3D topography on which we plotted the molecular images of biological specimen in an automated way with the ambition to make the system compatible with the use in the humans’ body. This is possible for two reasons. First, we have developed an automated system which moves the microprobe above the acquisition zone of the specimen and doesn’t require manual operation. This fact separates the Robot-Assisted SpiderMass from other MSI techniques. Second, the SpiderMass microprobe is mini-invasive and has already been demonstrated by the application on skin and *in vivo* during the canine patient surgery^3,7^. The system is envisioned for future clinical use in oncology surgery as an autonomous device, in contrast to simply coupling the probe to robot-assisted surgery devices, such as the MasSpec Pen to the Da Vinci robot^14^, where the robots still require manual operation by the clinicians. In this way, the automated robot-assisted system will move and screen the desired position indicated by the medical staff and obtained a real-time feedback that could be monitored directly by the medical team though augmented-reality by fusing the molecular data to the organ images. This type of system would leverage and ease the decision-making process during surgery. Taken together, we believe the intended developments and modifications of the modular robot-assisted system will forest the next generation of *in vivo* molecular image guidance used in a surgical room.

## METHODS

### Samples

Fresh-frozen Wistar rat brains were provided by the biology laboratory, University of Lille. The fresh-frozen brain was sectioned into coronal 12 μm sections using a microtome (Leica Biosystems, Wetzlar, Germany). The sponge piece and apple was purchased at the supermarket (Auchan, France). The human skin punch biopsy was purchased from C3(1)/SV40 T-antigen heterozygote transgenic mice were crossed, and breading pairs were tested for homozygosity. Negative mice were then used for this project. For post-mortem imaging a 3-4 month old male female mouse mice were was sacrificed, and post-mortem mice were immediately used for imaging.

### Ethical approval

Adult Wistar male rats (225-250g, 7-8 weeks old) were sacrificed according with European and french guidelines for animal research (European Convention for the Protection of Vertebrate Animals used for Experimental and other Scientific Purposes, ETS No.123) and approved by the local Animal Ethics committee (C2EA-075 Nord-Pas de Calais). Rats were maintained and housed under pathogen-free conditions at the University of Lille Animal Care Facility. All mice procedures were performed under a protocol (#01989.02) approved by the Animal Protocol Review Committees of the Institut Pasteur de Lille (France) in accordance with European regulation. C3(1)/SV40 T-antigen heterozygote transgenic mice colony is maintained and housed under pathogen-free conditions at the Institut Pasteur de Lille (France) Animal Care Facility.

### Hardware adjustments

The commercially available stiff 6D-axis precision MECA 500 robotic arm (MECADEMIC, Montreal, Canada) with repeatability of 5 μm, and less than 100 μm path accuracy. The distance sensor, composed of a camera (NIYPS Full HD 1080P) and a laser pointer, and the SpiderMass microprobe were attached to the robotic arm with a 3D-printed mount. The distance between each component was controlled and entered into a MATLAB code, to control the robot and acquire the topographical data and co-locate it with Spider Mass measurements.

### Distance Sensor

The sensor uses the angle of the camera to measure the distance: If the camera and the laser pointer are very close to the surface, the red point will be on the side of the recorded image. The higher the surface the closer the point will be to the center of the recorded image. In order to accurately measure the depth and cancel out the distortion effect of the camera lenses, the laser points are calibrated at different heights.

### Automatized user interface

The robotic arm and launching of other components (laser, distance sensor, camera) were controlled through the MATLAB coded framework. The distance sensor scanning and data collection were integrated into the code with manually setting the parameters such as height threshold, step size, scan zone area x,y; distance between the distance sensor and microprobe and z-height for optimal focal plane. To generate the topographical map, the distance sensor scanned all the x and y positions and registered all of the z-height differences. The x,y,z coordinates were exported as mesh to map for topographical visualization. After the topographical map was completed it was registered in the code and automatically launching the SpiderMass acquisition. The SpiderMass probe followed the same x,y,z topographical coordinates as registered in the scan. The SpidrMass laser was automatically firing by the artificial mouse clicking controlled by the Matlab Interface. The mass spectrometer was manually launched prior to the SpiderMass acquisition. The first intense chromatographic peak was indicative of the first pixel on the image.

### The SpiderMass system

The basic design of the instrument setup is described in detail elsewhere ^4,7^ In the following experiments, the prototype was equipped with a fibered tunable wavelength laser system between 2.8 μm to 3.1 μm (Radiant version 1.0.1, OPOTEK Inc., Carlsbad, USA) is pumped by the 1.064 μm radiation delivered by a Q-switched 10 ns pulse width Nd:YAG laser (Quantel Laser, Les Ulis, France). The IR laser microprobe is tuned to 2.94 μm to excite the most intense vibrational band of water (O-H). The major advantage of the SpiderMass system from other intraoperative mass spectrometry probes is its low invasiveness (few μm depth per shot according to the laser energy, 0.1-0.3 mm^3^ for a laser spot size of 400-500 μm^2^) and the ability to screen across the sampled tissue. A 1.5-meter length biocompatible laser fiber with 450 μm inner diameter (HP fiber, Infrared Fiber Systems, Silver Spring, USA) was connected to the exit of the OPO system and focused by a 20 mm focal length CaF_2_ lens attached at the end. A Tygon^®^ tubing (2.4 mm inner diameter, 4 mm inner diameter, Akron, USA) was used to aspirate the ablated material and is directly connected to the inlet of the mass spectrometer Xevo (G2-S, Q-TOF, Waters, Manchester, UK) through a modified atmospheric pressure interface described elsewhere^1^. The laser energy was fixed at 4 mJ/pulse corresponding to a laser fluence of 3.18J/ cm^2^ with variable number of laser shots.

### SpiderMass acquisition

The acquisition sequence was composed of 3 consecutive laser shots and 3 seconds between each step. The laser bursts were automatically fired by the MATLAB user interface. The data was acquired in positive and negative, sensitivity ion mode. The acquired spectra were averaged and scaled using MassLynx software (v4.2 SNC966, Waters Laboratory Informatics). MS/MS spectra were recorded after the isolation of precursor ion and they were subjected to collision-induced dissociation in the transfer cell with a transfer collision energy range between 25 to 40 V, depending on the selected precursor ion. The lipid annotations were performed manually through LIPIDMAPS database, Alex123 and METLIN database and the lipid assignment was realized with LIPIDMAPS and the literature^15–17^.

### Image processing

A second Matlab code was developed for topographical map visualization and image co-registration. Small topographical artifacts in the generated mesh were manually removed, if needed^8^. The raw mass spectrometry data was converted into mzxML file using an open source MS converter GUI^18^. The topographical map and mzxML file were imported into the MATLAB code. Several parameters such as the threshold of the chromatographic peak, the final time of acquisition, mass limits and height threshold were manually set. The selected mass limits were displayed on the topographical image. With the set parameters the calculated k value needed to correspond to the number of data points. Several image processing parameters such as normalization and transparency were accommodated in the code. In the current setup the absolute intensities of the mass spectrum were selected only from the local maximum of each chromatographic peak. The calculated pixel number should correspond to the topographical image matrix. The mass window is selected to display the distribution of the mass on the topographical image. The co-registered images were normalized to the TIC.

## ACKNOWLEDGMENTS

This work was funded by Ministère de l’Enseignement Supérieur, de la Recherche et de l’Innovation, Université de Lille and Inserm. The project was also funded by « Défi Santé numérique » joint Inserm/CNRS grant (I.F.) and ISite ULNE (Université Lille Nord-Europe) ERC Generator (I.F.).

## AUTHOR CONTRIBUTIONS

I.F. and N.O. wrote the manuscript original draft; A.K., C.D., I.F. and N.O. designed the experiments. N.O., P.C. and P.S. performed the experiments. C.L. took care of the mice models and assisted for the mouse experiments. I.F., N.O., P.C. and P.S analyzed the data. A.K., C.D., C.L., I.F., M.S., N.O., P.C and P.S. corrected the manuscript. C.D., A.K. and C.D. supervised the project. A.K., C.D., I.F. and M.S. provided the funding.

---

**Video 1. Topography MSI experiments inside of the body of the freshly sacrificed mouse.**

