## Supplementary Material for "Robot-Assisted SpiderMass for *in vivo* Real-Time Topography Mass Spectrometry Imaging"

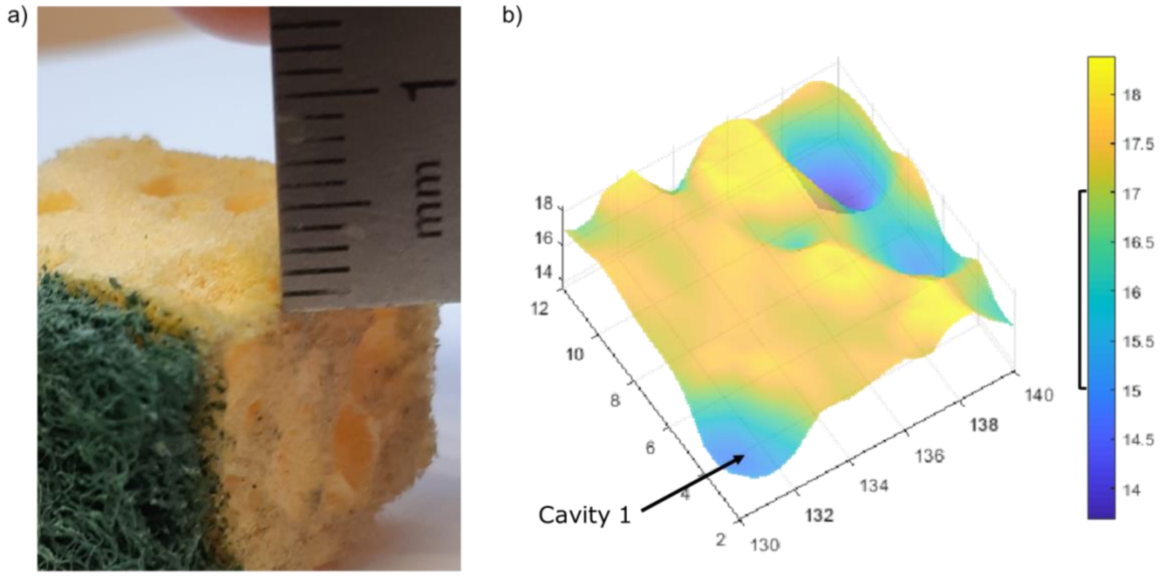


**Supplementary Figure 1. Topographical images of a sponge piece.** Images of cavity depths, **a)** measured by a ruler and **b)** reconstructed topographical image. The Cavity 1 is measured to be 2 mm deep by topography which is confirmed by measuring with the ruler.

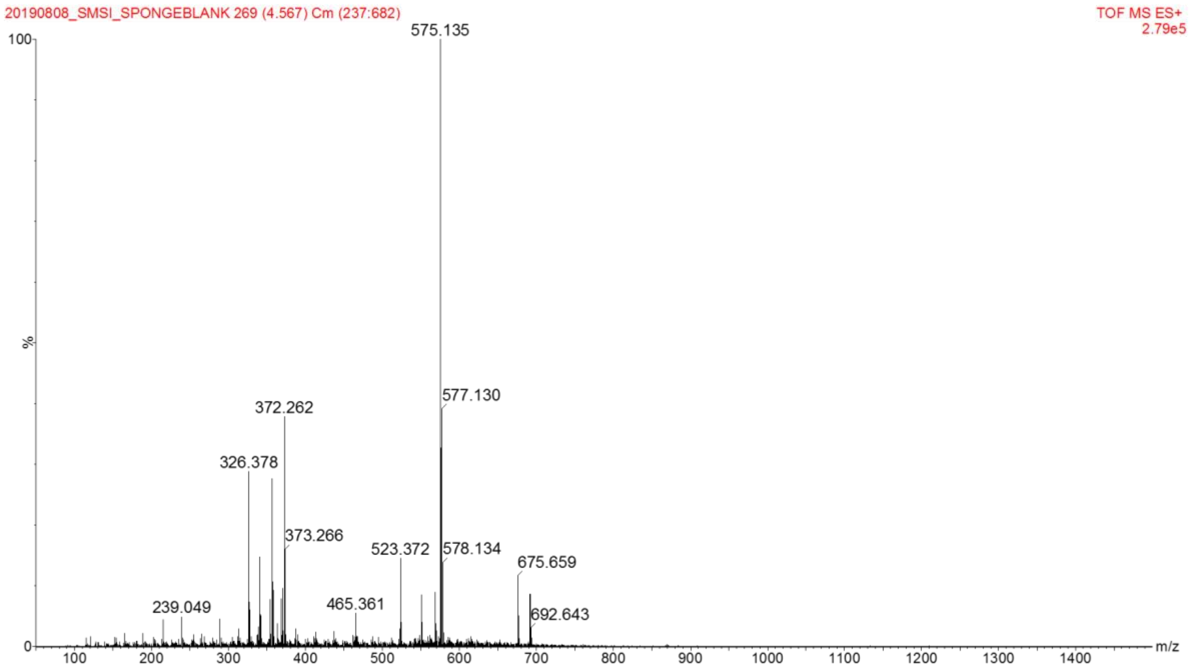


**Supplementary Figure 2.** SpiderMass mass spectrum of the sponge prior to lipid standard deposition measured in positive ion mode.

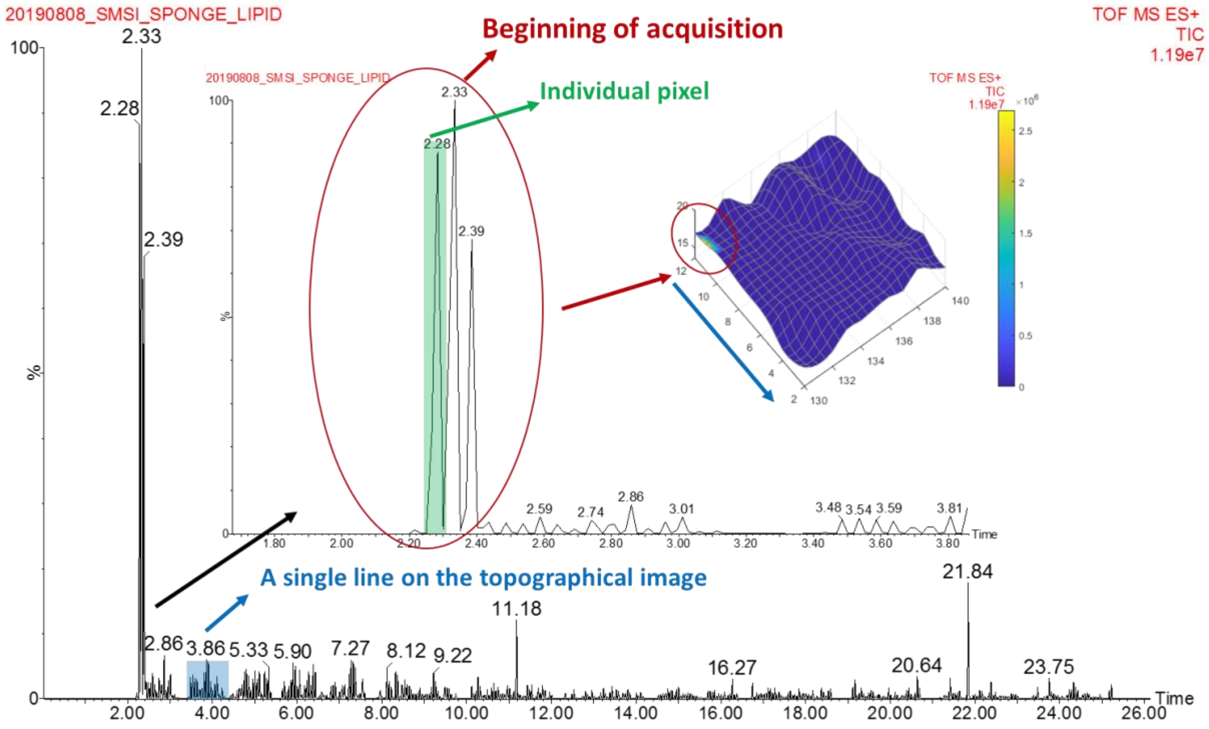


**Supplementary Figure 3. 3D-robotic topography imaging of a sponge piece.** TIC chromatogram of the acquired data across the 1cm$^2$ sponge surface following topographical image. The three major chromatographic peaks indicate the beginning of the acquisition and the second to fourth pixel on the reconstructed image. Each chromatographic peak corresponds to one pixel (green bar). Each group of peaks corresponds to a single line on the topographical image mesh (blue bar/blue arrow). The blue arrow shows the direction of the molecular data acquisition.

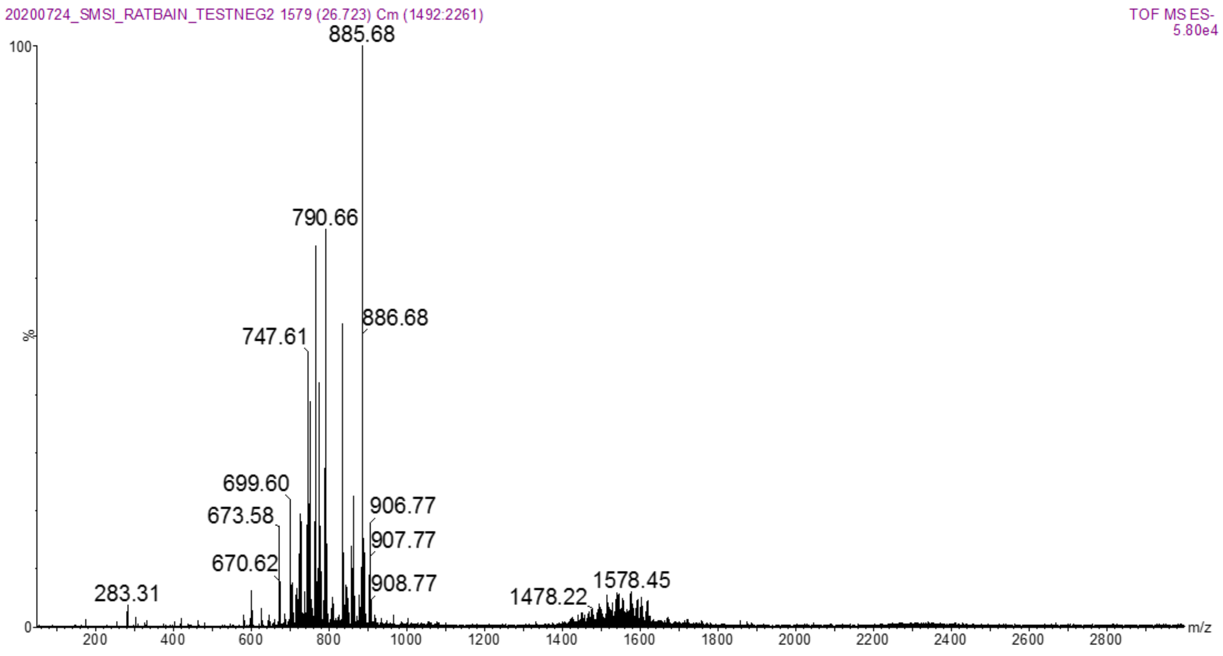


**Supplementary Figure 4. 2D-robotic assisted imaging of a rat brain section.** Averaged mass spectrum in the negative ion mode of the whole rat brain 2D image. The spectrum was scaled to the based peak (BP) and presented in relative intensity.

| Precursor ion | Detected ion fragments | Annotation |
|---|---|---|
| ***m/z* 726.57** | *m/z* 444.31 Loss of FA (18:1)<br>*m/z* 281.27 - FA (18:1)<br>*m/z* 196.05 ethanolamine headgroup | **[PE (18:1/18:1)-H]$^{-}$** |
| ***m/z* 747.6** | *m/z* 463.19 - Neutral loss of FA (18:0)<br>*m/z* 437.24 - Loss of FA (22:6)<br>*m/z* 327.21 - FA (22:6)<br>*m/z* 283.25 - FA (18:0)<br>*m/z* 152.98 - Glycerol-3-Phosphate (loss of $H_2O$) | **[PA (22:6_18:0)-H]$^{-}$** |
| ***m/z* 862.64** | *m/z* 826.72 – loss of HCl<br>*m/z* 383.38 – FA (26:6)<br>*m/z* 337.36 – FA (22:1)<br>*m/z* 283.28 – FA(18:0)<br>*m/z* 281.27 – FA (18:1) | **SHexCer d18:1;22:0***<br><br>**[PC(P-18:0_22:1))+Cl]$^{-*}$** |
| ***m/z* 885.65** | *m/z* 601.24 - Neutral loss of FA (18:0)<br>*m/z* 599.28 - Loss of FA (20:4)<br>*m/z* 437.24 - Loss of FA (20:4) and inositol<br>*m/z* 303.21 - FA (20:4)<br>*m/z* 283.25 - FA (18:0)<br>*m/z* 259.23 - Inositol phosphate ion<br>*m/z* 241.00 - Inositol phosphate ion (loss of $H_2O$)<br>*m/z* 152.98 - Glycerol-3-Phosphate (loss of H2O) | **[PI (18:0_20:4)-H]$^{-}$** |
| ***m/z* 890.67** | *m/z* 854.75 – loss of HCl<br>*m/z* 331.28 – FA(22:4)<br>*m/z* 309.30 – FA(20:1)<br>*m/z* 303.25 – FA(20:4)<br>*m/z* 283.28 – FA(18:0)<br>*m/z* 281.27 – FA(18:1)<br>*m/z* 255.25 – FA(16:0) | **SHexCer d18:1/24:0***<br><br>**[PC(O-42:2))+Cl]$^{-*}$** |

**Supplementary Table S1. Lipids identified from the 2D-robotic assisted imaging of a rat brain section.** MS/MS identification of the lipids observed during the imaging sequence from the rat brain tissue section in the negative ion mode. *Putative annotations when MSMS spectra were not sufficient for proper ID based on LIPID MAPS, Alex 123 and METLIN database.

| Precursor ion | Putative identification |
|---|---|
| ***m/z* 664.6** | $[M+H]^+$:HexCer 30:1;O2;<br>$[M+Na]^+$: Cer 40:1;O2;<br>$[M+NH_4]^+$: DG 37:6; Cer 40:1;O<br>$[M+K]^+$: Cer 39:2;O2; Cer 38:3;O3 |
| ***m/z* 700.6** | $[M+H]^+$: Cer 43:2;O3; PE O-34:3, Cer44:1;O2;<br>$[M+Na]^+$: PC O-29:0; PE O-32:0;<br>$[M+NH_4]^+$: DG 41:6; PA O-36:4;<br>$[M+K]^+$: Cer 43:2;O2; Cer 44:1;O |
| ***m/z* 737.6** | $[M+H]^+$: PG O-34:0; PA O-40:5; TG 43:0; PG 33:0; PA 39:5<br>$[M+Na]^+$: DG 43:4; TG 42:4; PA O-38:2; DG 44:11; PG dO-34:4; PA 37:2<br>$[M+NH_4]^+$: PE O-35:0; PC O-32:0; PC 31:0; PE 34:0; PE-NMe2 32:0; PS O-32:1; PS 31:1<br>$[M+K]^+$: CE 22:5 ; DG 42:5 |

**Supplementary Table S3. Putative ID of lipids from the 3D-robotic assisted imaging of the apple core.** Putative annotations where based on LIPID MAPS, Alex 123 and METLIN databases. The species were selected with $\Delta < 0.1$Da threshold.

| Precursor ion | Detected ion fragments | Annotation |
|---|---|---|
| **m/z 597.49** | MSMS inconclusive | **[DG (34:0)+H]$^{+*}$** **[DG 33:4+Na]$^{+*}$** |
| **m/z 701.6** | *m/z* 443.25– loss of FA(18:1)+$H_2O$ m/z 364.25- loss of FA (18:2) +$C_3H_6O_2$ m/z 337.25 – FA (18:2) +$C_3H_6O_2$ m/z 264.26 – Sphingosine+H-2$H_2O^+$ m/z 184.01 – PC headgroup | **SM 34:2;O2*** |
| **m/z 742.6** | *m/z* 683.48 – loss of $N(CH_3)_3$ *m/z* 601.52 – loss of PE headgroup *m/z* 443.25– loss of FA(18:1)+$H_2O$ m/z 337.25- FA (18:2) +$C_3H_6O_2$ m/z 184.01 – PC headgroup | **[PC-O-(34:3) +H]$^+$** **[PE (36:3) +H]$^+$** |
| **m/z 760.6** | m/z 557.01 – loss of PC headgroup m/z 522.32 – loss of FA(16:0) m/z 504.32- loss of FA(16:0)+$H_2O$ m/z 184.01 – PC headgroup | **[PC (34:1) +H]$^+$** |

**Supplementary Table S2. Lipids identified from the 3D-robotic assisted imaging of the skin biopsy.** MS/MS identification of the lipids observed during the imaging sequence from the disk skin biopsy in positive ion mode. *Putative annotations when MSMS spectra was not sufficient for proper ID based on LIPID MAPS, Alex 123 and METLIN database.

| Precursor ion | Detected ion fragments | Annotation |
|---|---|---|
| ***m/z* 616.19** | *m/z* 557.16 – loss of $CH_2COOH$<br>*m/z* 498.15 – loss of $CH_2COOH$ from the 557.16 ion | **Heme*** |
| ***m/z* 630.6** | m/z 612.6 – Cer+H-$2H_2O^+$<br>m/z 282.27 – Sphingosine+H-$H_2O^+$<br>m/z 264.26 – Sphingosine+H-$2H_2O^+$<br>m/z 252.27 - Sphingosine+H-$H_2O$-$HCHO^+$ | **Fragment of SM or GalCer**<br>**$[Cer + H\text{-}H_2 O]^+$** |
| **m/z 734.6** | *m/z* 551.51 – loss of PC headgroup<br>m/z 496.33 – loss of FA(16:0)<br>*m/z* 184.01 – PC headgroup | **$[PC (32:0)+H]^+$** |
| ***m/z* 834.6** | *m/z* 651.53 – loss of PC headgroup<br>*m/z* 578.53 –FA (18:0) -$H_2O$<br>*m/z* 551.51 – loss of FA (18:0)<br>*m/z* 184.01 - PC headgroup | **$[PC (40:6)+H]^+$** |
| ***m/z* 844.6** | *m/z* 785.45 – loss of $N(CH_3)_3$<br>*m/z* 611.52 – loss of PC headgroup<br>*m/z* 184.06 – PC headgroup<br>*m/z* 162.95 – cyclophosphane + $K^+$ | **$[PC(38:6)+K]^{+**}$** |
| ***m/z* 876.6** | m/z 603.53 -loss of FA (16:0)+$NH_3$<br>m/z 577.52- loss of FA (18:1)+$NH_3$<br>m/z 575.52- loss of FA (18:0)+$NH_3$<br>m/z 265.23 - RC=O- (18:1) | **$[TG (52:2) + NH_4]^+$** |

**Supplementary Table S3. Lipids identified from the 3D-robotic assisted imaging of the mouse.** MS/MS identification and tentative annotations of the lipids observed during the imaging sequence from the whole-body mouse experiment in the positive ion mode from the mammary gland, the heart and the lung. *Heme is already pre-charged and is thus not a protonated ion. **Putative annotations when MSMS spectra was not sufficient for proper ID based on LIPID MAPS, Alex 123 and METLIN database.
